# Trade-offs in biosensor optimization for dynamic pathway engineering

**DOI:** 10.1101/2021.04.20.440682

**Authors:** Babita K. Verma, Ahmad A. Mannan, Fuzhong Zhang, Diego A. Oyarzún

## Abstract

Recent progress in synthetic biology allows the construction of dynamic control circuits for metabolic engineering. This technology promises to overcome many challenges encountered in traditional pathway engineering, thanks to their ability to self-regulate gene expression in response to bioreactor perturbations. The central components in these control circuits are metabolite biosensors that read out pathway signals and actuate enzyme expression. However, the construction of metabolite biosensors is a major bottleneck for strain design, and a key challenge is to understand the relation between biosensor dose-response curves and pathway performance. Here we employ multiobjective optimization to quantify performance trade-offs that arise in the design and calibration of metabolite biosensors. Our approach reveals strategies for tuning dose-response curves along an optimal trade-off between production flux and the cost of an increased expression burden on the host. We explore properties of control architectures built in the literature, and identify their advantages and caveats in terms of performance and robustness to growth conditions and leaky promoters. We demonstrate the optimality of a control circuit for glucaric acid production in *Escherichia coli*, which has been shown to increase titer by 2.5-fold as compared to static designs. Our results lay the groundwork for the automated design of control circuits for pathway engineering, with applications in the food, energy and pharmaceutical sectors.

## I. INTRODUCTION

Metabolic engineering has led to a wealth of chemicals produced with microbial hosts. Classic pathway engineering employs combinations of heterologous expression and knock-downs of native enzymes of the host, so as to redirect metabolic flux toward production pathways^1,2^. Progress in synthetic biology and genetic engineering now allow the construction of control circuits that sense pathway activity and actuate enzyme expression in real time. These feedback systems are an attractive strategy to build pathways that self-tune to fermentation conditions and thus overcome many of the challenges encountered in pathway engineering. For example, it is generally challenging to find the right enzyme expression levels that maximise production and, at the same time, avoid excessive genetic burden on the host, flux bottlenecks, or the accumulation of toxic intermediates. Dynamic control circuits can resolve these challenges by self-adjusting enzyme expression in response to metabolic signals and continuously controlling pathway activity in response to intracellular or bioreactor perturbations.

Almost a decade ago, Zhang and colleagues reported the first successful implementation of such a control circuit, which was designed to improve production of fatty acid ethyl esters in *Escherichia coli*^3^. Since then there has been a sharp increase in the number of pathways and organisms where this technology has been deployed, and numerous reviews have discussed the opportunities and challenges offered by dynamic pathway engineering^2,4–8^. Control circuits have been built into diverse production pathways, including fatty acids^9^, carotenoids^10^, and other high-value chemicals^11,12^, using industrially-relevant hosts such as *Saccharomyces cerevisiae*^13,14^ and *Bacillus subtilis*^15,16^.

Current implementations of dynamic control differ substantially on their components and architectures, but generally speaking, control circuits require biosensors to read out metabolite concentrations and up-/down-regulate enzyme expression accordingly. Metabolite biosensors are thus central to dynamic pathway engineering, and as a result substantial efforts have been made toward their characterisation^17^. For example, a number of works have focused on the tuneability of metabolite-responsive transcription factors (TFs) using computational methods^18^ or large screens of biosensor dose-response curves^19,20^. Metabolite-responsive TFs have become the most widespread mechanisms for sensing because they can be tuned with many techniques, including promoter engineering^18,21^ and protein engineering^22,23^, and can be repurposed across species and pathways^24,25^.

Despite enormous progress in biosensor design, their construction is costly, labour intensive, and significantly slows down strain design. Biosensor dose-response curves depend on a complex interplay among various processes, such as metabolite-TF binding, TF binding to its target promoter site, and the expression dynamics of the TF itself. Such processes are poorly characterised in all but a handful of well studied TFs (e.g. the LacI repressor), and as a result biosensor design relies largely on iterative screens of construct libraries. Moreover, prior to implementation it is unclear how the shape of the biosensor dose-response curve affects production performance. In principle, such insights can be obtained from kinetic models based on ordinary differential equations (ODEs). Yet due to the inherent properties of metabolic systems, such as nonlinear kinetics, high-dimensionality and complex stoichiometry, it is generally challenging to analyse kinetic models with respect to the parameters of the biosensor doseresponse curve. Works in this space have primarily focused on the impact of biosensor parameters on metabolic flux using dynamical systems^26,27^, model simulation^28–30^, and control theory^31–33^. Other works have looked at how various control architectures, i.e. combinations of negative and positive feedback loops, affect the steady state response of engineered pathways, for example using exhaustive sampling of the design space^34^, or through bespoke mathematical analyses^35^. Most approaches employed so far, however, are *ad hoc* to specific pathways, and to date there are no generally applicable methods to study the relation between dose-response curves and pathway activity.

Here we present a method to identify performance tradeoffs in the design of metabolite biosensors across a wide range of metabolic engineering problems. Using multiobjective optimization^36–38^, we show that metabolite-responsive TFs can be designed to optimally trade-off competing objectives. Our focus is on the interplay between production flux and the costs associated with increased gene expression burden, a phenomenon known to affect cell physiology and impair production^39^. The method produces libraries of biosensors with varying dose-response parameters that optimally navigate the flux versus burden trade-off. We show the effectiveness of our approach in a simple model for a branched pathway that contains many features found in metabolic engineering problems, such as nonlinear kinetics, toxic intermediates, and a limited carbon supply. Using this exemplar system, we explore the performance trade-offs in three negative feedback architectures implemented in the literature^9,11,40^ and determined their robustness to fluctuations in carbon sources and leaky promoters.

We further tested our method in a more complex pathway for production of glucaric acid in *E. coli,* a high-value precursor for a number of applications. Based on published data, we built a detailed kinetic model for the pathway and computed optimal biosensors for different control architectures based on negative feedback. Our results show that a recently implemented control architecture^11^ outperforms alternative implementations of negative feedback, thus demonstrating the untapped potential of multiobjective optimization for dynamic pathway engineering.

## II. RESULTS

### A. Multiobjetive optimization of pathway dynamics

We focus on heterologous pathways described by kinetic models of the form:

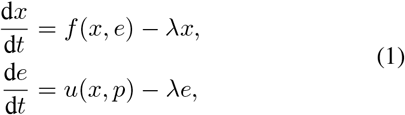

where *x* and e are vectors of metabolite and enzyme concentrations, respectively. The first-order term represents the dilution effect on molecular concentrations caused by cell growth with rate constant λ. The function *f* (*x*) describes the kinetic model in terms of mass balance equations, whereas the term *u*(*x,p*) is a vector-valued function that describes the expression rate for each heterologous enzyme, and p is a vector containing the parameters of the biosensors to be designed.

In the case of static pathways, heterologous enzymes are expressed at constitutive rates so that *u*(*x, p*) in Eq. (1) is constant and independent of pathway intermediates. Conversely, for pathways under dynamic control, the expression rates of pathway enzymes are made dependent on metabolite concentrations. In such case, each entry in *u*(*x,p*) describes the dose-response curve of a biosensor being employed to control heterologous expression. Different control architectures can thus be modelled via combinations of activation and repression feedback loops encoded in the shape of each doseresponse curve.

Here we adopt a multiobjective optimization approach to design the dose-response curves in *u*(*x,p*). As shown in Fig. 1A, in this setting dose-response curves correspond to points in a performance space, defined by the values of two objective functions to be jointly optimized. A large number of dose-response curves can be thus mapped onto a point cloud in a performance space shown in Fig. 1A. The boundary of such point cloud is the *Pareto front,* i.e. the curve containing all optimal trade-off designs where no objective function can be further reduced without increasing the value of the other^36^. Therefore, points along the Pareto front correspond to specific dose-response curves from which designers can choose biosensors to match a desired trade-off between the chosen performance objectives. For the purposes of biosensor design, the general optimization problem can be stated as:

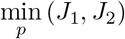

subject to:

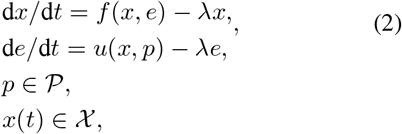

where *J*_1_ is inversely proportional to the production performance, and *J*_2_ quantifies the costs of pathway activity, as quantified e.g. by the amount of the heterologous enzymes expressed. The sets 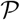 and *χ* represent constraints on the dose-response parameters and toxicity constraints on pathway metabolites, respectively. In Fig. 1B we illustrate the concept behind our approach. Starting from a kinetic model for a pathway of interest, plus a catalogue of candidate control architectures, our method produces libraries of biosensor dose-response curves that optimally navigate the performance space.

**FIG. 1.**
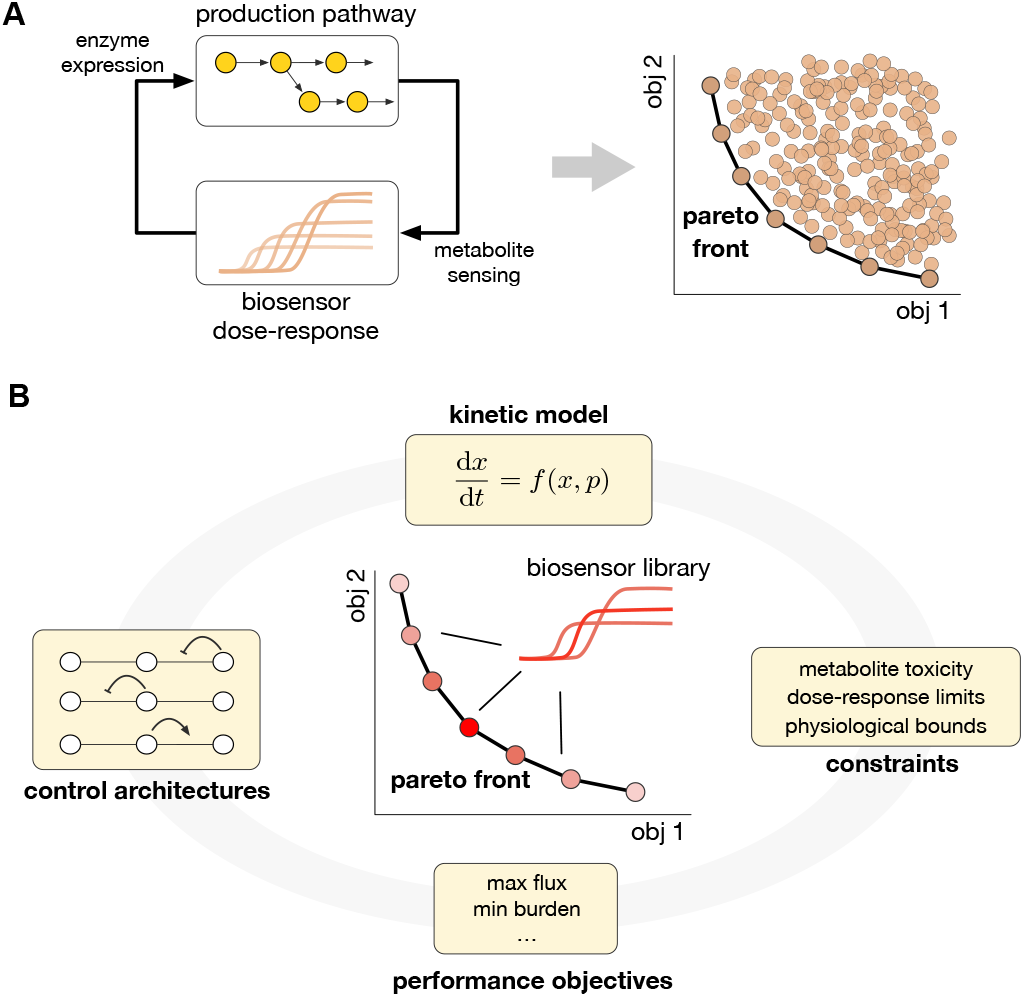
Performance trade-offs in engineered pathways with dynamic control. **(A)** General schematic for dynamic control in production pathways (left); a pathway intermediate is sensed by a metabolite-responsive transcription factor that controls the expression of pathway enzymes. Each biosensor dose-response curve can be mapped onto a point in a performance space (right). The outer boundary of the resulting point cloud is the *Pareto Front,* defined as the set of biosensors from which no objective can be further reduced without increasing the other^36^. **(B)** Design of biosensor libraries using multiobjective optimization. Starting from a kinetic model, biosensors for a specific control architecture can be optimized to trade-off performance objectives under various implementation constraints. Our approach addresses pathways in which the critical factors are the enzyme expression levels themselves, and thus excludes pathways with catalytically-compromised enzymes that require screening and protein engineering techniques for their optimization^41^.

The performance objectives in (2) can be chosen according to the specific application and pathway at hand, but generally they should represent the costs and benefits associated to pathway expression. In this paper, we focus on control circuits that weigh production flux against the increased expression load imposed by pathway enzymes. Production is often counteracted by the negative impact of heterologous expression on the physiology of the host. Pathway expression draws molecular resources away from endogenous processes, such as RNA polymerases, σ-factors, tRNAs and ribosomes. This phenomenon is generally known as “genetic burden”^42^ and can lead to homeostatic imbalances that impair growth and ultimately limit production^43,44^. This causes production flux and genetic burden to conflict with each other, in the sense that higher production flux entails a higher burden on the genetic machinery of the host.

### B. Open-loop optimization of a model pathway

To understand the baseline performance trade-offs in absence of dynamic control, we first carried out open-loop optimization of enzyme expression of a prototypical pathway that contains many features encountered in metabolic engineering problems. As shown in Fig. 2A, we consider the production of a metabolite *x*_1_ from a native intermediate *x*_0_. The two-step production pathway is catalyzed by heterologous enzymes *e*_1_ and *e*_2_. We model each reaction using Michaelis-Menten kinetics and assume that a carbon source is taken up at a constant rate *V*_in_; we have included the detailed model in the Methods section and all model parameters can be found in Supporting Data.

**FIG. 2.**
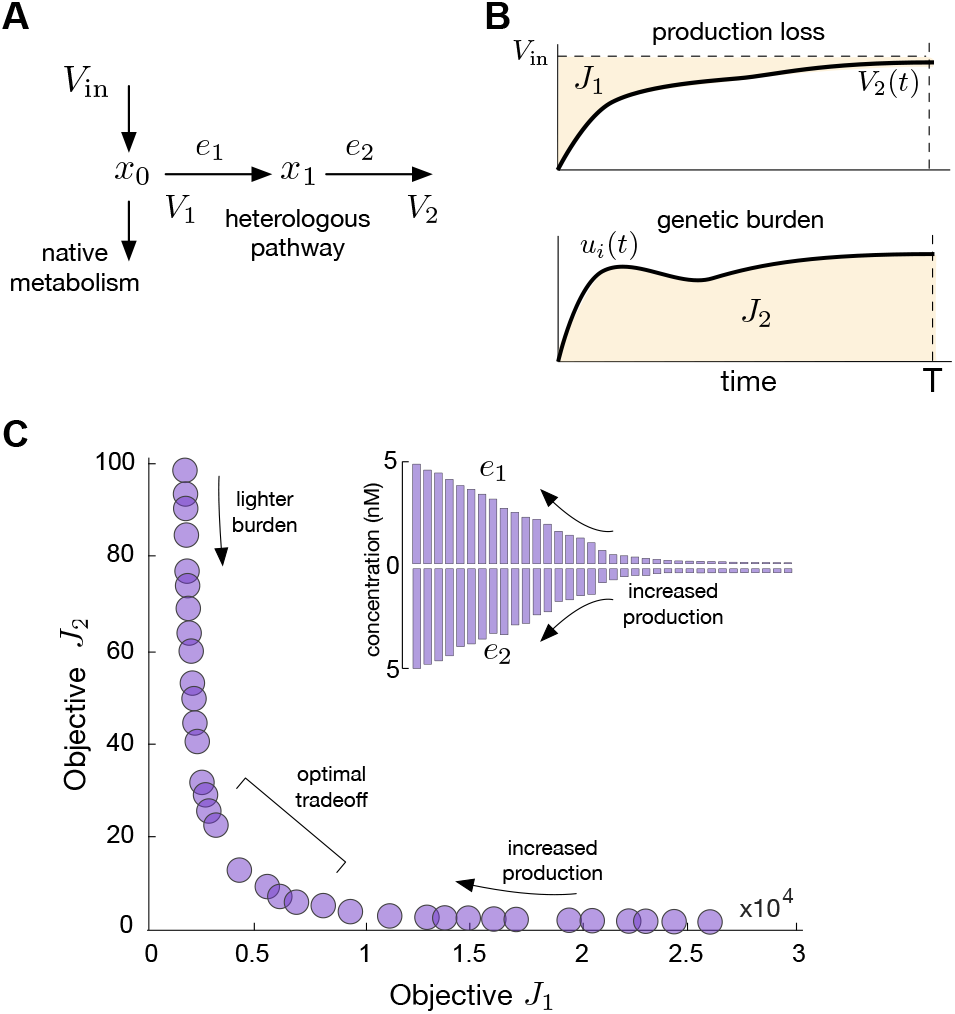
Exemplar model system. **(A)** Two-step engineered pathway branching out from central carbon metabolism. Carbon is taken up at a constant rate V_in_ and split between growth and the production of metabolite *x*_1_. **(B)** Objective functions employed in this paper for the optimization of biosensor dose-response curves, as defined in Eq. (4). **(C)** Pareto front of optimal solutions for the case of static control; insets show the optimal steady-state enzyme expression levels.

To gain an initial understanding of the design trade-offs of this model system, we first optimized its enzyme expression levels in open-loop, i.e. in the absence of dynamic control. This corresponds to the traditional approach in static pathway engineering, whereby heterologous enzymes are expressed at constant levels. In this case, the concentrations of both heterologous enzymes follow the ODE model:

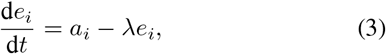

where *a_i_* is the rate of constitutive expression.

To model the trade-offs between production and burden, we define two objective functions to be minimised:

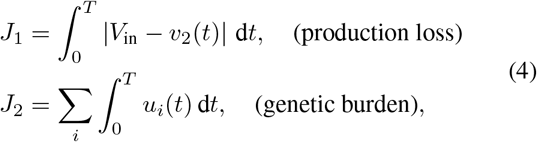

where *u_i_*(*t*) is the temporal expression rate of the *i*^th^ pathway enzyme, and *T* is an optimization horizon after activation of heterologous expression at *t* = 0. In the open-loop case, the enzyme expression rates are *u_i_*(*t*) = *a_i_* as in Eq. (3). The chosen objective functions, illustrated in Fig. 2B, quantify the production performance (*J*_1_) and the genetic burden (*J*_2_). The objective *J*_1_ describes the loss in pathway production, so that low values of *J*_1_ imply a high production flux *v*_2_(*t*) close to the carbon influx *V*_in_. The objective J_2_, conversely, corresponds to the total concentration of heterologous protein expressed during the temporal horizon. Note that for convenience, we have defined performance as a *production loss,* so that both objectives need to be jointly minimised.

We computed the enzyme expression rates *a_i_* as solutions to the multiobjective optimization problem of the form min (*J*_1_, *J*_2_); the full statement of the optimization problem is shown in Table IA. We imposed an upper bound on the maximal expression rates *a_i_* < 1 x 10^-3^μMs^-1^ to model the limited biosynthetic capacity of the host; this constraint corresponds to maximal enzyme concentrations of ~ 5μM in fast growing *E. coli* with doubling time of 26 min. We additionally introduced a constraint on the intermediate along the whole optimization horizon (*x*_1_(*t*) ≤ 180 μM), to model cytotoxic effects commonly encountered in pathway engineering^34^.

**TABLE I.**
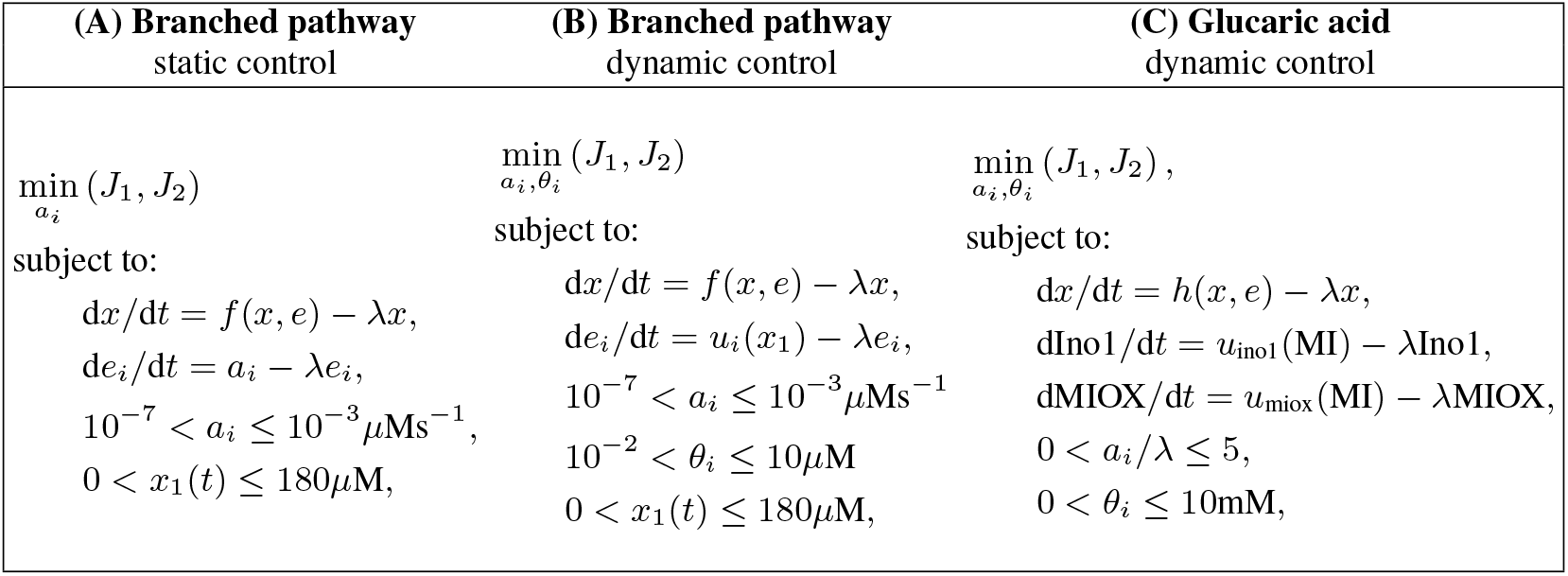
Multiobjetive optimization problems considered in this paper. In the first two cases, for the branched pathway in Fig. 2A the kinetic model *f* (*x*) is given in Eqs. (13)–(14). For the glucaric acid pathway in Fig. 5, we use the kinetic model *h*(*x*) defined in Eq. (S4). In each case, the enzyme expression rates *u_i_* are given in Eqs. (6) and (11). The full set of model parameters and optimal Pareto points be found in the Supporting Data.

| (A) Branched pathway<br>static control | (B) Branched pathway<br>dynamic control | (C) Glucaric acid<br>dynamic control |
| --- | --- | --- |
| $\min_{a_i} (J_1, J_2)$<br>subject to:<br>$dx/dt = f(x, e) - \lambda x,$<br>$de_i/dt = a_i - \lambda e_i,$<br>$10^{-7} < a_i \leq 10^{-3} \mu\text{Ms}^{-1},$<br>$0 < x_1(t) \leq 180 \mu\text{M},$ | $\min_{a_i, \theta_i} (J_1, J_2)$<br>subject to:<br>$dx/dt = f(x, e) - \lambda x,$<br>$de_i/dt = u_i(x_1) - \lambda e_i,$<br>$10^{-7} < a_i \leq 10^{-3} \mu\text{Ms}^{-1}$<br>$10^{-2} < \theta_i \leq 10 \mu\text{M}$<br>$0 < x_1(t) \leq 180 \mu\text{M},$ | $\min_{a_i, \theta_i} (J_1, J_2),$<br>subject to:<br>$dx/dt = h(x, e) - \lambda x,$<br>$d\text{Ino1}/dt = u_{\text{ino1}}(\text{MI}) - \lambda \text{Ino1},$<br>$d\text{MIOX}/dt = u_{\text{miox}}(\text{MI}) - \lambda \text{MIOX},$<br>$0 < a_i/\lambda \leq 5,$<br>$0 < \theta_i \leq 10 \text{mM},$ |

We solved the optimization problem using a genetic algorithm detailed in the Methods section; all solutions can be found in Supporting Data. The resulting Pareto front, shown in Fig. 2C, has three distinct regimes: (i) low production and light burden, (ii) high production and heavy burden, and (iii) an optimal trade-off regime between the two. These regimes are in agreement with the intuition that stronger enzyme expression levels (shown in insets of Fig. 2C) lead to higher production flux at the expense of an increased cost in protein expression. The convexity of the Pareto front indicates that the optimization problem is well-posed, in the sense that both objective functions oppose each other across the whole space of optimal expression levels.

### C. Optimal biosensors for negative feedback control

We next turned our attention to the design of dynamic control circuits for the general branched pathway of previous section. As shown in Fig. 3A, we consider three control architectures in which the biosensor detects the concentration of intermediate (*x*_1_) and controls enzyme expression accordingly. For generality, we assume that the biosensor is implemented via a metabolite-responsive transcription factor (TF), which is currently the most widespread strategy for implementation of metabolite biosensors^2^.

**FIG. 3.**
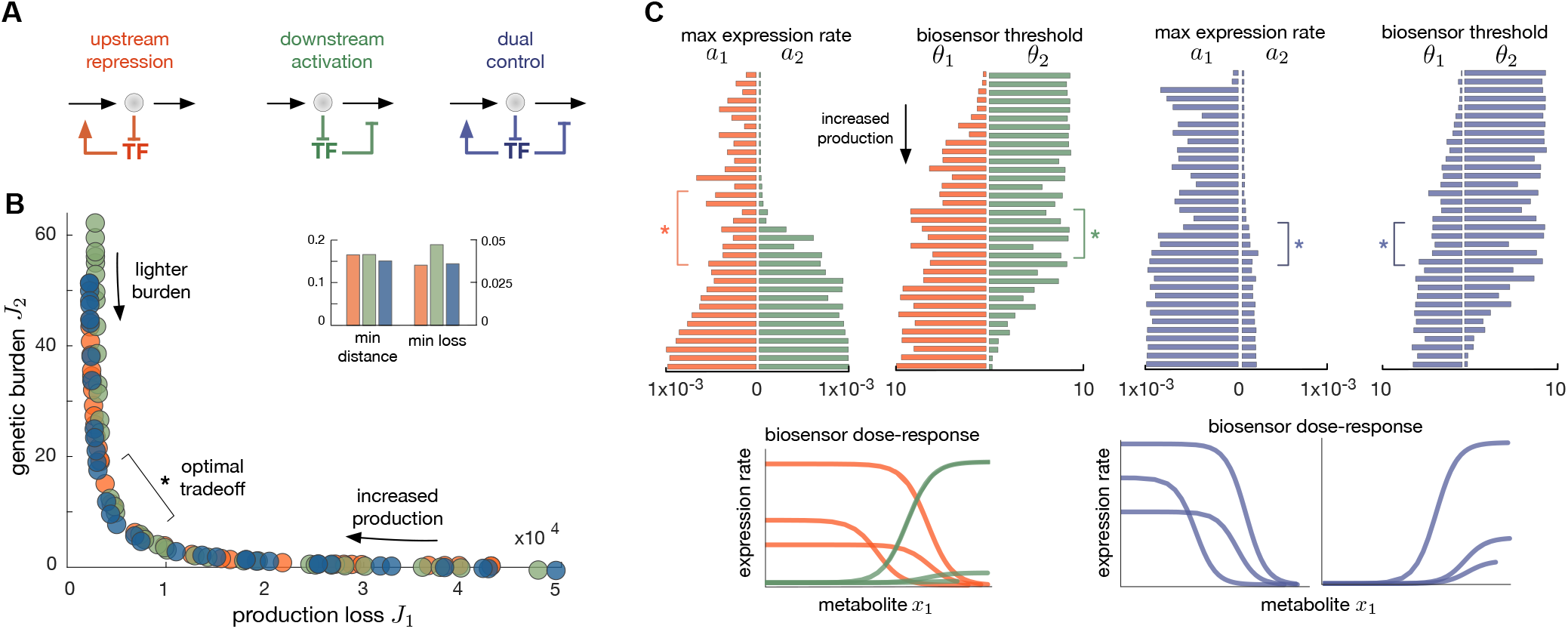
Optimal bionsensors in a model pathway. **(A)** We consider three negative feedback architectures for the model pathway in Fig. 2A. In all cases, feedback is implemented through a metabolite-responsive TF. **(B)** Pareto fronts of the considered negative feedback circuits. The inset shows the minimum distance to the origin for each front, as well as the minimal production loss in each case; both metrics were computed from a 10^th^ order log-log polynomial regression for the Pareto front normalised to the [0,1] range in both coordinates. These results were computed for a fixed value of the effective Hill coefficient in all cases (*h_i_* = 2); as shown in Supplementary Figure 1, we also found that the Pareto fronts are largely insensitive to the chosen Hill coefficient. **(C)** Optimal biosensor libraries for the considered control architectures; the bars represent the optimal dose-response parameters (maximal expression rate *a_i_* in μMs^-1^, threshold *θ_i_* in μM) for each optimal biosensor along the Pareto fronts in panel B. Bottom insets show three representative dose-response curves for each circuit.

In this case, as in Eq. (1) the expression of heterologous enzymes follows the ODE model:

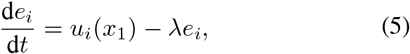

where *u_i_*(*x*_1_) is the dose-response curve of the biosensor with respect to the the concentration of the intermediate. In this case, there are a total 3^2^ −1 = 8 possible control architectures, because each of the two enzymes can be subject to three modes of control (positive, negative, and no control) and we exclude the open-loop case studied in the previous section. In this work we focus on biosensors that act on the basis of molecular sequestration, whereby binding between the TF and the intermediate x1 causes a reduction in the pool of regulators that can bind to the operator region of enzymatic genes. This mechanism produces a form of negative regulation and is found in various biosensors employed in the literature so far, including BetI^45^, FadR^3^, IpsA^11^, FapR^9^, and LacI^23^. Other relevant sensing mechanisms, which are not included by our modelling approach, are co-repressors such as TrpR that bind to DNA only upon metabolite binding^46^, and LysR-type transcriptional regulators such as BenM^47^ that bind constitutively to different DNA domains depending on the presence or absences of a metabolite.

As shown in Fig. 3A, we focused on three different implementations of negative feedback: a transcriptional repressor that downregulates upstream enzymes (upstream repression), a transcriptional activator that upregulates downstream enzymes (downstream activation), or a TF with dual regulatory function that combines both control strategies (dual control). We deliberately excluded circuits that include positive feedback loops, such as upstream activation or downstream repression, as these display multistable dynamics that are generally not desirable in production pathways, with the exception of particular use cases^35,48^. Negative feedback is also known to offer various benefits, including improved robustness^26^, accelerated response times^40^, and noise control^35^. The three considered architectures have been implemented in several metabolic control systems in *E. coli*. For example, an upstream repression circuit was employed for speeding up the production of fatty acids using the regulator FadR as biosensor^40^. The downstream activation circuit has been recently built for production of glucaric acid with IpsA as a metabolite-responsive biosensor^11^. The dual control architecture was employed for the production of malonyl-CoA using FapR as a dual transcriptional regulator^9^.

By optimizing the biosensor dose-response curves *u_i_*(*x*_1_) for each control architecture, we computed libraries of biosensors that optimally navigate the production-burden space. To this end, we modelled the dose-response curves as sigmoids:

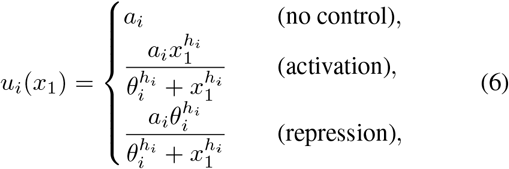

where *a_i_* is the maximal expression rate proportional to the promoter strength and expression level of the TF, *θ_i_* is a regulatory threshold, and *h_i_* is an effective Hill coefficient to model cooperative effects. Note that the sigmoidal doseresponse curves in Eq. (6) are effective models that lump together various molecular processes, including metabolite-TF and TF-DNA binding. For design purposes, such lumped models are preferred because they are simpler to parameterise with biosensor characterization data^18^, as compared to full kinetic models of TF expression and binding. Moreover, implementations of dynamic control systems often put the TF under constitutive expression^2,9^, thus reducing the impact of TF expression dynamics on pathway performance.

In applications, the effective Hill coefficient *h_i_* is particularly challenging to tune experimentally and thus we fixed it to *h_i_* = 2 in all our examples, focusing instead on optimizing parameters *a_i_* and *θ_i_*, which are more readily accessible with standard promoter engineering techniques. To optimize the biosensor parameters *α_i_* and *θ_i_*, for each architecture we solved a multiobjective optimization problem of the form min (*J*_1_,*J*_2_) subject to constraints on maximal enzyme expression (*α_i_*), biosensor thresholds (*θ_i_*), and a toxicity constraint on the intermediate xi as in the previous case. The full statement of the optimization problem is shown in Table IB, and all solutions are given in Supporting Data.

As shown in Fig. 3B, to quantitatively compare the Pareto fronts of each circuit we computed their minimum distance to the origin (indicative of the optimal trade-off achieved by each architecture) and the minimal production loss (representing the ability to maximise the production flux). Overall, we observe that the three circuits achieve similar Pareto fronts with a nearly identical optimal trade-off. Yet we found that the minimal production loss is ~30%. higher in the downstream activation circuit, which means that the best production achievable by downstream activation is substantially worse than the other two circuits. Taken together, the analysis suggests that upstream repression and dual control are similarly strong in terms of performance. However, this also suggests that the extra cost of implementing the more complex dual control system does not offer advantages in terms of the trade-offs considered in this paper.

Detailed inspection of the biosensor libraries reveals that architectures require different designs to optimize performance (Fig. 3C). For example, both upstream repression and downstream activation require increasing promoter strengths (*a_i_*) to increase production. Yet the optimal promoter regulatory thresholds (*θ_i_*) exhibit opposite trends. In the upstream repression circuit, increasing the regulatory threshold helps to increase production, but the downstream activation circuit requires a lowered threshold to increase production. In the case of the dual control architecture (Fig. 3C), we observe a strong asymmetry between promoters. The upstream promoter needs to be at least 5-fold stronger than the downstream one, and their regulatory thresholds follow opposite trends, similarly as in the first two architectures. Altogether these results suggest that strategies for fine-tuning the trade-off between production and burden are highly dependent on the chosen control architecture.

### D. Robustness and sensitivity of the control circuits

The optimization results in Fig. 3A suggest that the Pareto fronts of the three negative feedback circuits are similar to the open-loop case in Fig. 2C. While this seems counter-intuitive, from control theory it is known that open-loop control can indeed outperform feedback systems, at the expense of their robustness to perturbations^49^. Open-loop solutions are fragile in the sense that their performance can significantly degrade in face of perturbations or inaccuracies in the model of the system to be controlled. We hence sought to identify salient properties of the circuits in terms of their robustness to perturbations commonly encountered in applications: a) fluctuations in growth conditions, and b) leaky expression of promoters.

#### a. Robustness to growth conditions

To compare architectures in terms of their robustness to carbon supply, we challenged the optimal circuits with perturbations to the influx *V*_in_. We first simulated each optimal circuit to steady state under a constant *V*_in_, and then introduced a step perturbation of a given size. As shown in Fig. 4A, we computed the relation between changes to *V*_in_ and the resulting change in production flux *V*_2_ for each circuit architecture and each combination of optimized parameters. Ideally, the production flux should be insensitive to growth conditions; this would translate into a flat curve in Δ*V*_2_ vs. Δ*V*_in_ in Fig. 4A. At the other end, fragile designs would produce a 45° line in the Δ*V*_2_ vs. Δ*V*_in_ space, meaning that the circuit is completely unable to compensate for perturbations in carbon influx. The latter case is what happens in static pathways, whereby carbon flux perturbations are relayed directly onto the flux of production pathways. In contrast, by sensing changes in the concentration of the intermediate (*x*_1_), the feedback circuits adjust enzyme expression so as to keep the production flux close to its level prior to the perturbation.

**FIG. 4.**
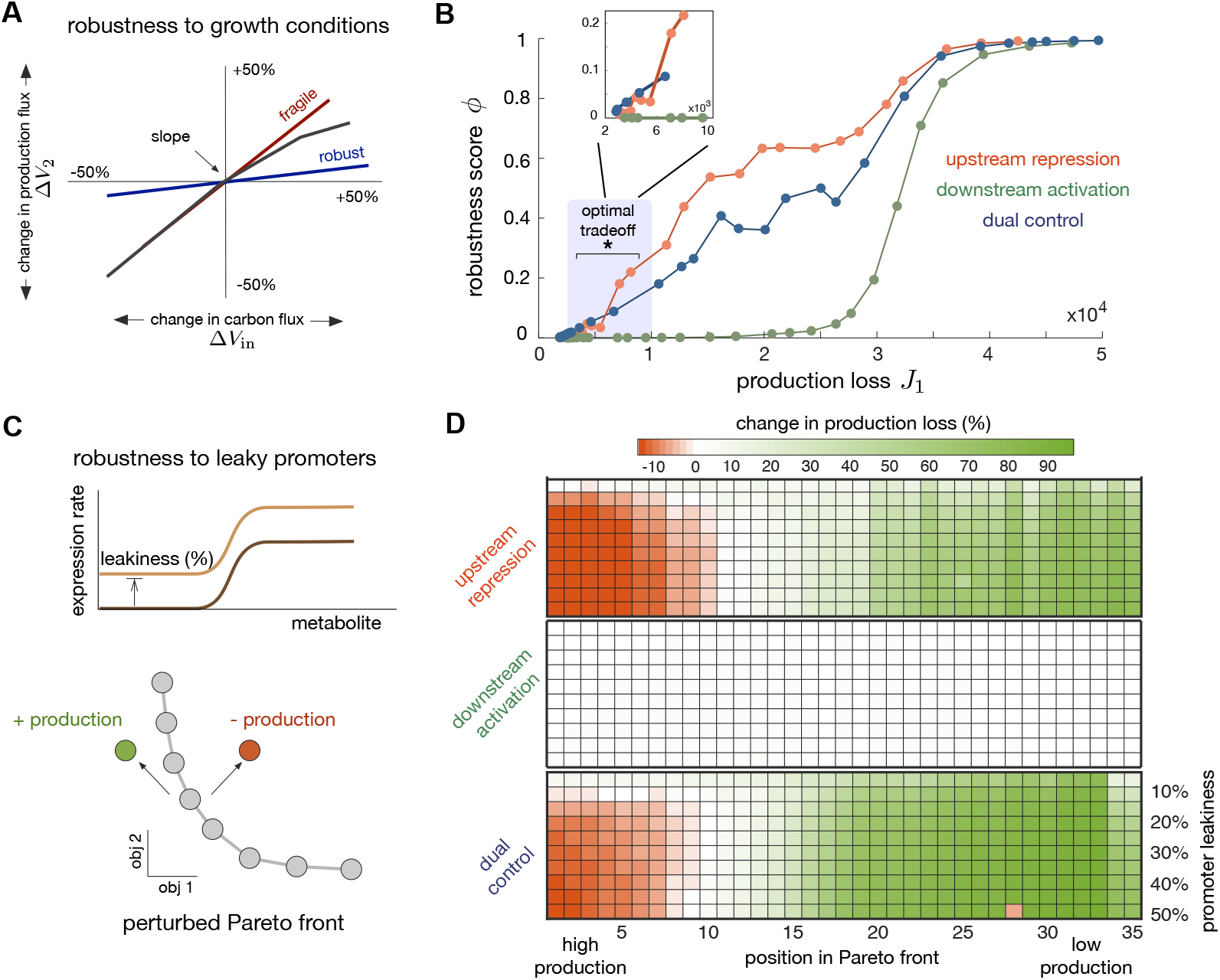
Robustness of optimal control circuits. **(A)** Robustness to changes in carbon flux. For each Pareto optimal design and each architecture in Fig. 3B, we computed the relative change in steady state production flux (Δ*V*_2_) as a function of a relative change in carbon influx (*V*_in_). We use the slope at zero of the resulting Δ*V*_2_ vs. Δ*V*_in_ curves as a metric for robustness. Circuits with a near nil slope are more robust than those with a slope close to unity. We first simulated a constant carbon influx *V*_in_ = 1μMs^-1^ until steady state and then introduced a step change in *V*_in_ until a new steady state is reached. The black line represents a typical response to perturbation in carbon influx; full simulation results for all circuits can be found in Supplementary Figure 2. **(B)** Robustness score *ϕ* =1 — slope as defined in (7) for all optimal designs in Fig. 3B. We found that upstream repression outperforms others and that downstream activation is particularly fragile to growth conditions. Zoom of optimal trade-off region shown in inset plot. **(C)** Robustness to leaky promoters. We simulated each optimal circuit in Fig. 3B with a modified model for gene expression that includes promoter leakiness, as shown in Eq. (8). From these simulations we computed the negative change in production loss Δ*J*_1_, defined in Eq. (9). The bottom panel illustrates the two ways in which leakiness can shift the Pareto points. **(D)** Heatmaps show the negative change in production loss (Δ*J*_1_) defined in Eq. (9) when circuits are simulated with promoter leakiness; shown are the three circuits and all optimal designs along the Pareto front in Fig. 3B. Upstream repression and dual control display regimes with lowered production (red, Δ*J*_1_ < 0), despite the additional enzyme expression caused by leaky promoters; this phenomenon appears in all tested leakiness levels, modelled by *a* in Eq. (8). The downstream activation circuit, in contrast, appears to be extremely robust to leaky expression, with Δ *J*_1_ < 2% across all designs and leakiness levels.

We observed substantial variations in the response of the three control circuits to perturbations in the carbon influx (see Supplementary Figure 2). For example, the downstream activation circuit fails to compensate for drops in carbon influx, possibly due to its resemblance with coherent feed-forward loops that are known to respond differently to positive and negative perturbations^50^. In contrast, the other two architectures display a compensatory response across both negative and positive perturbations. To quantitatively explore such differential responses, we defined the robustness score:

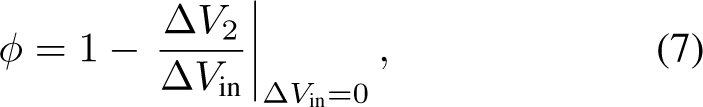

where 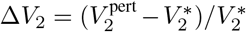 and 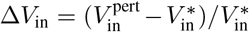 are relative changes in both steady state fluxes, and 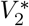 is the optimal steady state production flux achieved with a constant 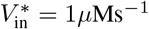. The second term in Eq. (7) is the slope of the Δ*V*_2_ vs. Δ*V*_in_ curves when Δ*V*_in_ = 0, as marked in Fig. 4A; all the computed curves for each control circuits can be found in Supplementary Figure 2.

Under the *ϕ* metric, robust circuits would score *ϕ* ≈ 1 while fragile designs would score as *ϕ* ≈ 0. We thus computed the robustness metric for every optimal design along the three Pareto fronts in Fig. 3B. The results, shown in Fig. 4B, suggest an inverse relation between production and robustness to carbon influx; the three circuits display poor robustness (*ϕ* ≈ 0) in the high production regime of the Pareto front, and high robustness (*ϕ* ≈ 1) in the low production regime. Moreover, we found that upstream repression outperforms the other architecture in terms of robustness to growth conditions, and this happens even in the optimal trade-off area of the Pareto front (highlighted in Fig. 4B). The downstream activation circuit appears to be particularly fragile, with near zero *ϕ* score across large regions of its Pareto front, except at the low production regime. Dual control, on the other hand, displays an intermediate level of robustness across many production levels, but as in the previous section, given its extra complexity and cost, it does not provide obvious benefits over upstream repression.

#### b. Sensitivity to promoter leakiness

Regulated promoters typically express a residual amount of protein in the absence of a regulator. This is commonly known as leakiness and is generally detrimental for circuit design. Since the mechanisms behind leaky expression depend on the specific regulatory mechanism of the promoter, leakiness is poorly understood and makes it difficult to predict system performance.

To quantify the dependencies between leakiness and optimal performance, we re-simulated each circuit in Fig. 3C with a modified model for enzyme expression:

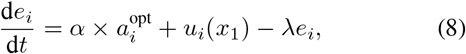

where 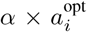 represents the promoter leakiness as a fraction of the optimized parameter 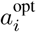, and *u_i_*(*x*_1_) is the doseresponse curve of the biosensor assuming a non leaky promoter. To quantify the robustness of each architecture to the leakiness a, we computed the change in the production loss:

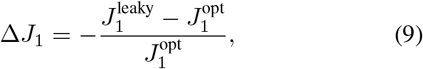

where 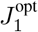 is the optimal production loss computed for non-leaky promoters, and 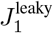 is the value of the production loss obtained with a leaky promoter. Note that for convenience we have defined Δ*J*_1_ as the negative change in loss. Intuitively, since leakiness increases the total amount of enzyme expressed, we expect a higher genetic burden accompanied by an increase in production, i.e. a decrease in the production loss *J*_1_ and Δ*J*_1_ > 0. On the contrary, designs where leakiness causes a drop in production would lead to Δ*J*_1_ < 0, an undesirable scenario where the extra genetic burden is accompanied by a lower production flux. This concept is illustrated in Fig. 4C, where perturbations to the Pareto front can shift designs to the left (more production, Δ*J*_1_ > 0) or to the right (less production, Δ*J*_1_ < 0).

We found stark differences in the Δ*J*_1_ scores for each architecture (Fig. 4D). Both upstream repression and dual control display regimes with lowered production flux (Δ*J*_1_ < 0), despite the additional availability of enzymes caused by leaky expression. Moreover, this phenomenon appears in the high production regime in both cases and across all tested levels of leakiness. This suggests that leaky expression leads to sub-optimal production in architectures that include an upstream repression component. Upstream repression can prevent an increase in the production flux due to the interplay between an increased enzyme abundance and the stronger repression caused by elevated concentration of the intermediate metabolite. In contrast, we found the downstream architecture displays negligible Δ*J*_1_ scores across all optimal designs and all tested levels of leakiness; the highest value for downstream activation in Fig. 4D is Δ*J*_2_ ≈ 2%. This result suggests that downstream activation produces a pathway flux that is extremely robust to promoter leakiness.

### E. Control circuits for production of glucaric acid

To illustrate how our approach can be used in a realistic metabolic engineering problem, here we applied the method to production of glucaric acid in *Escherichia coli*. Glucaric acid is a high-value precursor to several key products such as polymers^51^ and therapeutics^52^. The work by Doong an colleagues^11^ successfully built a dynamic control circuit that resulted in a 2.5-fold increase of glucaric acid titer compared to static designs. We thus used our approach to quantify the optimal performance of their circuit as compared to other alternatives that could have been implemented with similar genetic parts.

The seminal work of Moon *et al*^53^ built a pathway that synthesises glucaric acid from glucose-6-phosphate (g6p) in upper glycolysis. As shown in Fig. 5A, the four-step pathway requires the native *E. coli* enzyme inositol monophosphatase (SuhB), plus three heterologous enzymes: inositol-3-phosphate synthase (Ino1) from *Saccharomyces cerevisiae*, myo-inositol oxygenase (MIOX) from *Mus musculus,* and uronate dehydrogenase (Udh) from *Pseudomonas syringae*.

**FIG. 5.**
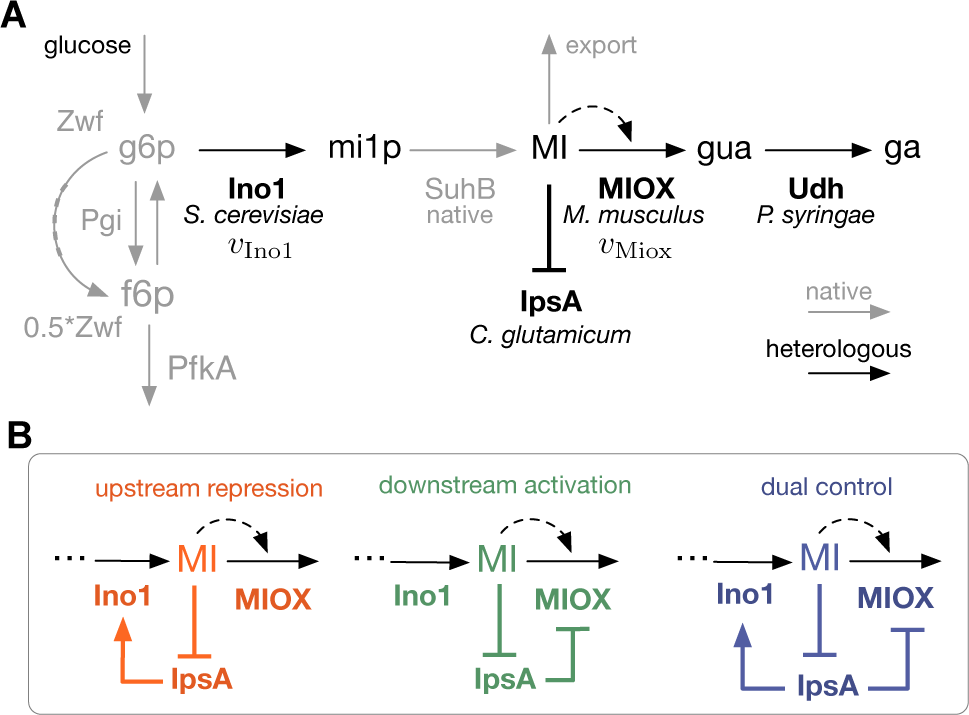
Pathway for synthesis of glucaric acid in *Escherichia coli.* **(A)** Synthetic pathway for production of glucaric acid from g6p in central metabolism of *E. coli^53^*. **(B)** Architectures for dynamic control using the *myo*-inositol-responsive transcriptional regulator IpsA. The downstream activation circuit has been implemented by Doong *et al*^11^.

In a subsequent work^11^, a dynamic control circuit was built using the HTH-type transcription factor IpsA, a dual transcriptional regulator from *Corynebacterium glutamicum* that is sequestered by pathway intermediate *myo*-inositol (MI). The circuit was engineered to put MIOX under negative control by IpsA, thus creating a negative feedback system akin to the downstream activation circuit in Fig. 3A. We employed our multiobjective optimization framework to compare the architecture implemented by Doong *et al^11^* against alternative negative feedback architectures, exploiting IpsA as a dual transcriptional regulator to explore the three architectures as in the previous exemplar model: downstream activation of MIOX, upstream repression of Ino1, and dual control of both Ino1 and MIOX (Fig. 5B).

We built a kinetic model of the glucaric acid pathway in Fig. 5A with parameters and enzyme kinetics mined from the literature and calibrated to multi-omics data^54^. To develop a minimal model, we assume that the reactions catalyzed by SuhB and Udh occur spontaneously, so pathway dynamics can be accurately captured by a reduced model for Ino1 and MIOX alone, as shown in Fig. 5B. This assumption is based on the observation that myo-inositol-1-phosphate (mi1p) and glucuronic acid (GUA) were not detected in culture products^53^, indicating that SuhB and Udh are not rate limiting. Moreover, activity of Udh is also reported to be two orders of magnitude faster than than Ino1^53^, providing further support that Udh is not rate limiting. The model also includes the reported leakage of *myo*-inositol to the media^53^, as well as the allosteric activation of MIOX by its own substrate MI (dashed black arrow in Fig. 5B). Further details of model equations, assumptions and parameter values are given in the Methods and the Supporting Information.

We modelled the Ino1 and MIOX reactions in Fig. 5B using Michael-Menten kinetics parameterised with *K*_m_ and *v*_max_ values sourced from literature (see Supporting Data). Since the enzyme turnover rates (*k*_cat_) were not available, we opted to model enzyme expression as fold-change factors that scale the *v*_max_ value. This strategy is based on the observation that maximal reaction rates scale linearly with enzyme concentration *E*, i.e. *v*_max_ = *k*_cat_ x *E*. We thus write the maximal rates as 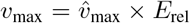, where 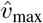 is the value from literature and *E*_rel_ is a non-dimensional variable representing relative enzyme abundance. We model the dynamics of the relative abundances of Ino1 and MIOX through the ODEs:

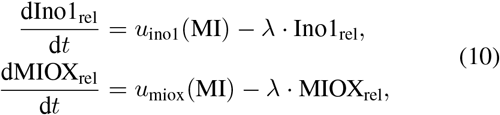

where the expression rates follow a lumped model:

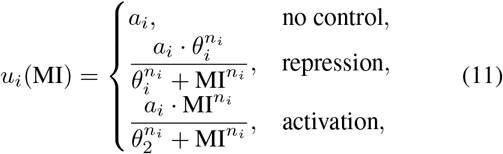

with *i* = {ino1, miox}. In this model, *a_i_* has units of s^-1^ representing the maximal expression rate and MI is the concentration of *myo*-inositol sensed by IpsA. The parameters to be optimized are *a_i_* and regulatory thresholds (*θ_i_*), while we fix the Hill coefficients to *n*_ino1_ = *n*_miox_ = 1. As in the previous example, our aim is to determine control designs that maximise production whilst minimising the burden of expressing Ino1 and MIOX. To this end, we define two objective functions to be minimised:

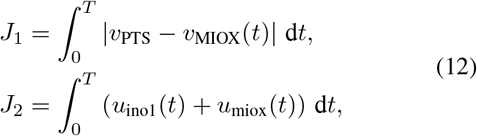

where *T* is an optimization horizon chosen as 10 times the doubling time log 2/λ. We formulated a multiobjective optimization problem to jointly minimise both objectives *J*_1_ and *J*_2_, as in the previous example. We additionally imposed bounds on the maximal expression rates (*a_i_*), as well as upper limits on the biosensor thresholds (*θ_i_*). The full statement of the optimization problem can be found in Table IC and all solutions are reported in the Supporting Data.

The optimization results in Fig. 6A indicate that both objective functions mutually oppose each other and that the Pareto front is convex in all cases. However, unlike the previous example, we found that downstream activation outperforms the other two architectures: for the same level of genetic burden, the downstream activation circuit produces a higher production flux. This difference can be attributed to the pathway structure, as the glucaric acid pathway includes a number of processes absent from the general branched pathway of the previous section, including allosteric substrate activation of MIOX, export of *myo*-inositol, and a more detailed model of central carbon metabolism. Similarly to the results in Fig. 3C, here we observe that higher MIOX expression generally improves production for the downstream activation architecture. The optimal steady state expression levels shown in Fig. 6C suggest that production can be increased with higher Ino1 expression, reflected by the linear increase in optimal Ino1 expression, and low expression of MIOX. Increases in Ino1 expression drive a higher accumulation of *myo*-inositol that results in a higher flux through the MIOX reaction. Examination of the MIOX turnover rate *v*_MIOX_/MIOX (i.e. the reaction per unit enzyme), shown in Fig. 6D, suggests that biosensors in the optimal trade-off region of the Pareto front tend to maximise MIOX turnover rate. Moreover, comparison of Fig. 6C-D indicates that Ino1 and MIOX levels have opposite effects on the turnover rate. Although increases in Ino1 boost the efficiency of the MIOX reaction, increasing MIOX expression itself has the converse effect and leads to lower efficiency. This is because higher amounts of MIOX reduce the steady state pool of *myo*-inositol, which in turn drives MIOX away from saturation. As a result, in the Pareto front a higher level of MIOX is needed to counteract the low turnover rate, causing the pronounced increase in burden and marginal gains in production observed at the top-left of the Pareto front in Fig. 6A. This observations are also true of the other feedback circuits (Supplementary Figure 3, Supplementary Figure 4). Altogether, these observations reiterate the complex dependencies between system performance and design parameters. These results provide strong computational evidence that the circuit architecture implemented by Doong *et al^11^* has substantial benefits over other implementations of negative feedback control.

**FIG. 6.**
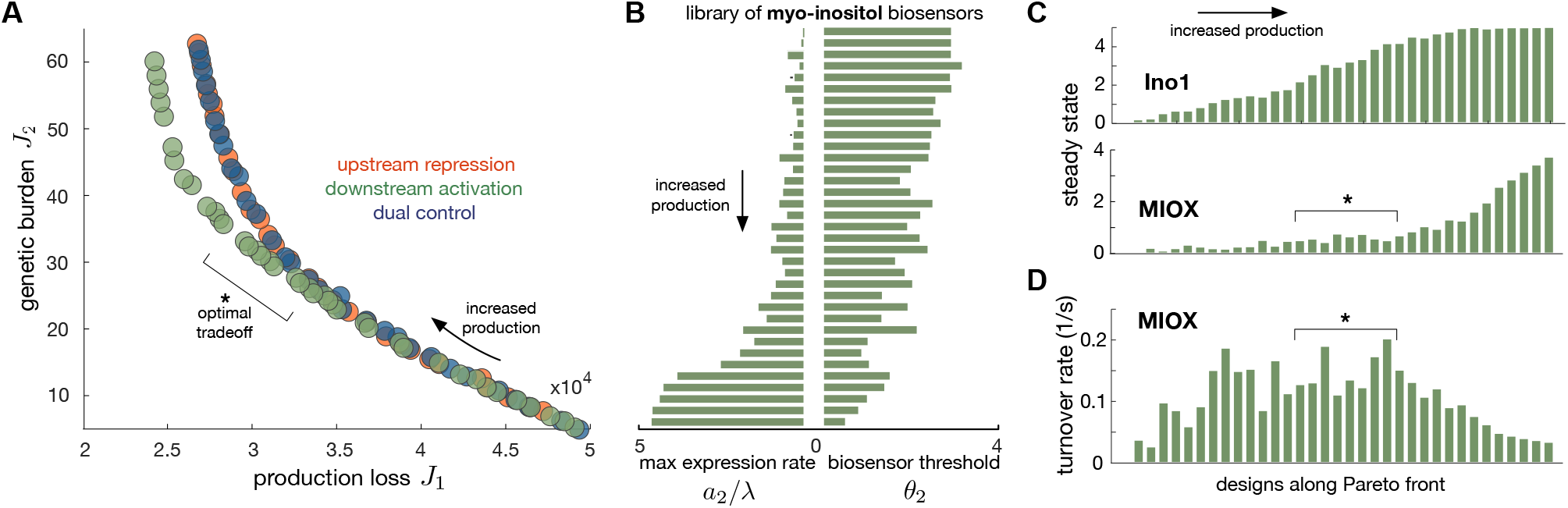
Multiobjective optimization of a glucaric acid production under dynamic control. **(A)** Pareto front for each control architecture in Fig. 5B; the downstream activation architecture implemented by Doong *et al^11^* outperforms the other two control strategies. **(B)** Optimal library of myo-inositol biosensors; bars represent the maximal expression rate in dimensionless units (*a*_2_/λ) and threshold (*θ*_2_) for the downstream activation architecture. Optimal solutions for upstream repression and dual control can be found in Supporting Data. **(C)** Steady state foldchange in abundance of Ino1 and MIOX enzymes for each optimal solution along the Pareto front of the downstream activation circuit in panel A. **(D)** Steady state turnover rate (rate per unit enzyme) of MIOX for each optimal solution in along the Pareto front of the downstream activation circuit.

## III. DISCUSSION

Dynamic control has emerged as a promising solution to common challenges in metabolic engineering, such as the optimization of enzyme expression timing and strength, avoidance of metabolic intermediate accumulation, and suboptimal performance in large fermentations. Previous dynamic control systems have been built mostly on the basis of an intuitive understanding of pathway features. In general it is extremely challenging to identify suitable control architectures and parameters via purely experimental trial-and-error approaches. Although computational methods promise to accelerate design and prototyping in synthetic biology applications, currently there are no such methods speficically tai-loref for dynamic pathway engineering. This is in stark contrast with static pathway design, where designers can avail of powerful computational methods^55,56^ and a large collection of software tools for analysis and design^57^.

The application of optimization methods to understand dynamics of natural pathways has a long history^58^. For example, the seminal work by Zaslaver et al^59^ showed that gene regulation of amino acid biosynthesis in *E. coli* could be understood as the solution of a cost-benefit optimization problem. Subsequent work^60–62^ validated some of those early observations in other organisms, demonstrating that optimization can be a powerful tool to reverse-engineer control systems found in nature. The focus, however, has been almost exclusively on natural systems and in the case of engineered pathways, the literature has largely centred on *ad hoc* simulation-based methods, resulting in a lack of general design methodologies that can be employed in a wide range of pathways.

Here we have shown that multiobjective optimization can be a powerful tool for dynamic pathway engineering. We considered several example control circuits and pathways, including various negative feedback circuits built in the literature, and benchmarked their optimal performance in terms of production flux and genetic burden on their host. The analysis revealed a number of trade-offs in biosensor optimization, indicating that different implementations of negative feedback control can display substantially different performance and robustness properties. For example, we considered a prototypical pathway (Fig. 2A) in which an upstream repression architecture appears to be beneficial in terms of performance and robustness to growth conditions, yet fragile to promoter leakiness. Conversely, downstream activation architectures have a poorer performance but are extremely robust to promoter leakiness. Such trade-offs have been studied extensively in control engineering, where it is known that optimal performance is often accompanied by poor robustness to perturbations^63^, yet their characterization for dynamic pathway control remain largely unexplored in the literature. Although it would be highly desirable to uncover general performance and robustness properties of specific control circuits, such as those discovered for natural metabolic pathways^60–62^, our results suggest that such design trade-offs are highly dependent on the pathway stoichiometry, enzyme kinetic parameters, and the specific metrics employed to quantify performance and robustness. This makes it challenging to establish general “design principles” for dynamic pathway control and highlight the utility of our method as a systematic approach to identify trade-offs for specific pathways and candidate control circuits selected for implementation.

To illustrate how our approach can be employed in real metabolic engineering tasks, we applied the optimization strategy to the production of glucaric acid in *E. coli*. This is an excellent model system to test our approach because other groups have already developed successful dynamic control strategies for it, and it contains a number additional complexities in its stoichiometry and post-translational regulatory mechanisms. Our approach predicts that the downstream activation circuit implemented by Doong et al^11^ substantially outperforms other negative feedback architectures. Each of the considered architectures require different genetic parts and constructs, and thus carry different costs in terms of practical implementation. The use of optimization-based designed thus allows to single out specific architectures that provide superior performance. Moreover, these results overall suggest that optimal pathway performance, and the control circuits that achieve it, are both highly specific to the pathway of interest. This highlights the utility of model-based approaches to biosensor design, as such complex dependencies between biosensor parameters, control architectures, and performance are impossible to detect from intuition alone.

In its most general formulation, our approach allows the integration of design objectives and constraints into an optimization problem that can be numerically solved with algorithms available in standard scientific computing software. The optimal solutions represent the biosensor dose-response curves that optimally navigate the performance space specified by the objective functions. In this work we have deliberately focused on genetic burden (cost) and production flux (benefit) as objective functions, on the basis that they are relevant across a wide range of design scenarios. However, the use of our framework in specific applications may require bespoke objective functions that capture known bottlenecks and limitations of the target pathway. For example, other relevant objectives in applications are the temporal concentrations of metabolites, robustness, or cell-to-cell heterogeneity^64,65^.

Similarly, here we focused on constraints on the parameters of the dose-response curve and upper bounds on toxic metabolites, as these are commonly encountered in applications. Optimization-based design also allows the inclusion of whole range of other relevant constraints, including upper bounds on enzyme concentrations, biophysical bounds on promoter parameters^66^, or thermodynamic limits for metabolic fluxes^67^.

We envision a number of promising extensions of our approach. First, in our models we have generally assumed that a constant growth rate in our models. A growing body of work^42,68–71^ has shown that genetic burden is the result of competition for cellular resources between native and heterologous genes, ultimately affecting growth rate^43^. Although to date there is no standardised or widely adopted strategy to model such burden effects, their inclusion would provide more insights into the design trade-offs normally encountered in applications. A key challenge is that current burden models, such as small lumped models of gene expression^69,71^ or large mechanistic models for growth^68^, require large amounts of data for parameterization that are typically not available in metabolic engineering applications. Second, for simplicity we have focused on metabolite biosensors implemented via sequestration-based TFs, because they are the most widely used mechanism used in dynamic pathway engineering. Our methodology is directly applicable to systems in which the biosensor can be well described by sigmoidal dose-response curves. But a number of works have successfully employed alternative sensing mechanisms for pathway control, e.g. transcriptional co-repression^72^, nucleic acid aptamers^73,74^, or metabolite-sensing riboswitches^75,76^. These mechanisms can be incorporated into optimization approaches, but this will require bespoke models different from the ones employed in our work. Other strategies, such as the use of quorum sensing to use culture density as a proxy for growth rate^77,78^, or the use of gene regulatory circuits to gain finer control of gene expression^79,80^ also require bespoke models once their implementation becomes more standardised^7^. As the repertoire of sensing mechanisms and control strategies grows, we expect that model-based design will become increasingly important for the identification of key design parameters and their impact on pathway performance.

Dynamic pathway engineering is significantly more complex than traditional static designs, because control circuits have more degrees of freedom and their effectiveness varies with pathway structures. A key step for the widespread adoption of this technology is the development of systematic design methods that make dynamic control more widely and easily adoptable. Here we have proposed a first step into such a strategy that shows promise across a range of metabolic engineering problems.

## IV. METHODS

### A. Computational models

#### a. Branched pathway

The mass balance equations for the exemplar pathway (Fig. 2A) in continuous culture are:

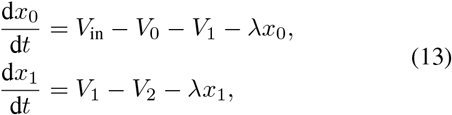

where *x*_0_ and *x*_1_ are the metabolite concentrations, and λ is the culture growth rate fixed at λ = 1.93 x 10^-4^s^-1^ (corresponding to a fast doubling rate of 26min in *E. coli*). The reaction rates are assumed to follow Michealis-Menten kinetics:

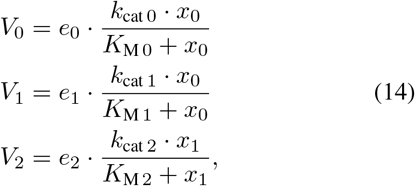

where *e_i_* are the enzyme concentrations and (*k*_cat *i*_, *K*_M *i*_) are kinetics parameters of each enzyme; we used representative values for the kinetic parameters^81^ and for simplicity assumed equal kinetics for the three enzymes. The model further assumes constitutive expression of the native enzyme *e*_0_, and regulated expression of the heterologous enzymes *e*_1_ and *e*_2_, as described in Eq. (6). All kinetic parameters can be found in the Supporting Data.

#### b. Glucaric acid pathway

The mass balance equations for the GA pathway (Fig. 5A) in continuous culture are:

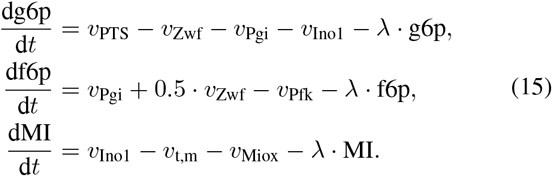

We assume a constant uptake of glucose *v*_PTS_ and growth rate λ, with values *v*_PTS_ = 1.34 mmol/gDCW/h = 0.1656mM/s and λ = 0.1 h^-1^ = 0.278 x 10^-4^ s^-1^ taken from the Keio multi-omics data set^54^. Details on the model construction, assumptions, reaction kinetics and parameters can be found in the Supporting Information and Supporting Data.

### B. Model simulation and optimization

In all cases we first simulated pathway dynamics in absence of heterologous enzymes, and use the resulting steady state metabolite concentrations as initial conditions for the dynamic control simulations. This strategy emulates common lab protocols in which strains are first pre-cultured in absence of heterologous expression, and then inoculated into fresh media containing inducers of enzyme expression^11^. The initial conditions for each case study can be found in the Supporting Data.

The full statement of the optimization problems solved in Fig. 2, Fig. 3, and Fig. 6 can be found in Table I. All numerical calculations were done in Matlab 2019b. The model for the branched pathway was integrated with the `odel5s` stiff solver, and the glucaric acid model was integrated with the Matlab implementation of the SUNDIALS `CVode` solver for precision and speed^82^. In all cases, parameter optimization was done with the multiobjective optimizer `gamultiobj` with parameter bounds and path constraints to account for cytotoxic effects of the intermediate. In all analyses, we computed a total of 70 Pareto points. For ease of visualisation, in Fig. 2, Fig. 3, and Fig. 6 we plotted every other solution, i.e. 35 out of 70 solutions. All optimized dose-response parameters are reported in the Supporting Data.

## SUPPORTING INFORMATION

Supporting Information contains details of model construction for myo-inositol pathway, plus Supporting Figures and the full result of the optimization algorithms

## ACKNOWLEDGEMENTS

BKV was funded by a Wellcome Trust ISSF3 fund awarded to DAO. AAM acknowledges funding from BBSRC Research Grant BB/M017982/1.

## AUTHOR CONTRIBUTIONS

BKV constructed and analyzed the model of the branched pathway; AAM constructed and analyzed the model of the glucaric acid pathway. DAO designed and supervised the research; all authors contributed to analysis and discussion of results.

