## Supplementary Information for "Trade-offs in biosensor optimization for dynamic pathway engineering"

### KINETIC MODEL FOR THE GLUCARIC ACID PATHWAY

Here we detail the construction of the kinetic model of the glucaric acid synthesis pathway in *E. coli* based on the construction in the work of Moon *et al*[53], shown in Fig. S1A. The pathway is composed of the product of a native gene *suhB* and three heterologous genes, *ino1* from *S. cerevisiae*, *miox* from *M. musculus* and *udh* from *P. syringae*.

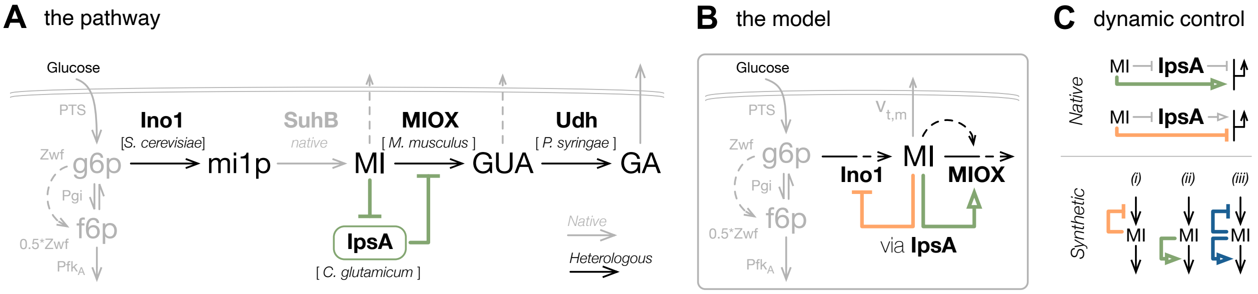


FIG. S1. **Glucaric acid synthesis pathway and its dynamic control.** (a). The pathway illustrated as constructed in *E. coli*[53]. Native enzymes are highlighted in gray and heterologous enzymes in black. The *myo*-inositol (MI)-dependent dynamic control via TF IpsA as engineered in[11] is shown in green. (b) Illustration of our reduced model of the metabolic pathways and their dynamic control. Reactions SuhB and Udh are assumed to occur spontaneously, so rate limiting steps Ino1 and MIOX capture the pathway dynamics. (c) We exploit IpsA as a dual transcriptional-regulator in its native host and design MI-activating and MI-inhibiting expression (top) to study three negative feedback architectures: (i) MI-inhibition of *ino1*, (ii) MI-activation of *miox*, and (iii) dual control.

We model the synthesis of glucaric acid in continuous culture. To this end, we consider a constant but limited availability of glucose, at a fixed rate, and a fixed culture growth rate. To capture this in our model, we define a fixed glucose uptake rate ($v_{\text{PTS}}$) and a fixed growth rate ($\lambda$) consistent with that uptake rate as:

$$v_{\text{PTS}} = 1.34 \text{ mmol/gDCW/h} = 0.1656 \text{ mM/s}, \tag{S1}$$

$$\lambda = 0.1 \text{ h}^{-1} = 0.278 \times 10^{-4} \text{ s}^{-1}, \tag{S2}$$

sourced from Keio multi-omics dataset[54]. To ensure that we simulate the model for long enough for the enzymes to reach steady state, we define optimization horizon $T$ as ten-fold the cell doubling time:

$$T = 10 \cdot \frac{\log 2}{\lambda} = 69.3 \text{ h} = 2.50 \times 10^5 \text{ s}, \tag{S3}$$

where $\lambda$ is defined in Eq. (S1). Our kinetic model is based on the following assumptions:

i) *Reactions SuhB and Udh occur spontaneously*

It has been reported that no *myo*-inositol-1-phosphate or glucuronic acid was detected in culture products[53], which indicated that SuhB and Udh are not rate limiting reactions. Moreover, Udh specific activity is reportedly two-orders of magnitude higher than Ino1 or MIOX[53], adding support that Udh is not rate limiting. As a result, we model the conversion of g6p directly into *myo*-inositol as a single reaction with the kinetics of Ino1. Likewise, we model the conversion of *myo*-inositol directly into glucaric acid as a single reaction with the kinetics of MIOX. The simplified model is shown in Fig. S1B.

ii) *Myo-inositol leaks into media*

*Myo*-inositol was detected in the culture products of the glucaric acid producing strain[53]. Moreover, its secretion was observed against the concentration gradient[83], indicating its active transport. We incorporate this leakage in the model as the reaction rate $v_{\text{t,m}}$ in Fig. S1B. (Table S1)

iii) *Substrate activation of MIOX activity*

Supplementing high concentrations of *myo*-inositol into the media enhanced activity of MIOX between 5–15-fold[53]. We posited that MIOX is allosterically activated by MI. This is modelled by rate equation $v_{\text{Miox}}$, with kinetic parameters estimated from specific activity data reported in[53].

iv) *Use of SUMO-MIOX fusion protein instead of MIOX* MIOX without the SUMO fusion tag degrades quickly so cannot retain long term production. SUMO-MIOX is more stable and we parameterize its kinetics as the average activity measured over 48 hours after the first 24 hours, as reported in[83].

v) *Rapid equilibrium of MI-IpsA complex*

We model expression of Ino1 and MIOX directly as a function of *myo*-inositol. By assuming rapid kinetics of *myo*-inositol binding with IpsA, we can phenomenologically model the expression dynamics of IpsA controlled enzymes as a Hill function dependent only on *myo*-inositol; such lumped models are standard in the literature because the parameters of the binding kinetics are difficult to measure[18].

#### Dynamics of pathway metabolites

Under the above assumptions, the mass balance equations for the pathway metabolites (g6p, f6p, and MI in Fig. S1B) are:

$$\begin{aligned}
\frac{\text{dg6p}}{\text{d}t} &= v_{\text{PTS}} - v_{\text{Zwf}} - v_{\text{Pgi}} - v_{\text{Ino1}} - \lambda \cdot \text{g6p}, \\
\frac{\text{df6p}}{\text{d}t} &= v_{\text{Pgi}} + 0.5 \cdot v_{\text{Zwf}} - v_{\text{Pfk}} - \lambda \cdot \text{f6p}, \\
\frac{\text{dMI}}{\text{d}t} &= v_{\text{Ino1}} - v_{\text{t,m}} - v_{\text{Miox}} - \lambda \cdot \text{MI}.
\end{aligned} \tag{S4}$$

TABLE S1. **Reaction rates for the kinetic model in Fig. S1B.**

| Reaction | Mechanism | Reference |
|---|---|---|
| $v_{\mathrm{Pgi}} = \frac{v_{\mathrm{m,pgi}} \cdot (\mathrm{g6p} - (\mathrm{f6p}/K_{\mathrm{eq,pgi}}))}{\mathrm{g6p} + (K_{\mathrm{m,pgi,g6p}} \cdot (1 + (\mathrm{f6p}/K_{\mathrm{m,pgi,f6p}})))}$ | Reversible Michaelis-Menten | [84] |
| $v_{\mathrm{Zwf}} = \frac{v_{\mathrm{m,zwf}} \cdot \mathrm{g6p}}{K_{\mathrm{m,zwf,g6p}} + \mathrm{g6p}}$ | Michaelis-Menten | [85] |
| $v_{\mathrm{Pfk}} = \frac{v_{\mathrm{m,pfk}} \cdot \mathrm{f6p}^{n_{\mathrm{pfk}}}}{K_{\mathrm{m,pfk,f6p}}^{n_{\mathrm{pfk}}} + \mathrm{f6p}^{n_{\mathrm{pfk}}}}$ | Hill equation | [84] |
| $v_{\mathrm{Ino1}} = \mathrm{Ino1}_{\mathrm{rel}} \cdot \frac{v_{\mathrm{m,ino1}} \cdot \mathrm{g6p}}{K_{\mathrm{m,ino1,g6p}} + \mathrm{g6p}}$ | Michaelis-Menten | [85] |
| $v_{\mathrm{t,m}} = \frac{v_{\mathrm{m,t,mi}} \cdot \mathrm{MI}}{K_{\mathrm{m,t,mi}} + \mathrm{MI}}$ | Michaelis-Menten | [53] |
| $v_{\mathrm{Miox}} = \mathrm{MIOX}_{\mathrm{rel}} \cdot \frac{v_{\mathrm{m,miox,eff}} \cdot \mathrm{MI}}{K_{\mathrm{m,miox,mi}} + \mathrm{MI}}$<br>where $v_{\mathrm{m,miox,eff}} = v_{\mathrm{m,miox}} \cdot \left(1 + \frac{a_{\mathrm{miox}} \cdot \mathrm{MI}}{K_{\mathrm{a,miox,mi}} + \mathrm{MI}}\right)$ | Michaelis-Menten<br>with substrate activation | [83] |

TABLE S2. **Parameter values for the kinetics in Table S1.**

| Param | $v_{\mathrm{pgi}}$ | $K_{\mathrm{eq,pgi}}$ | $K_{\mathrm{m,pgi,g6p}}$ | $K_{\mathrm{m,pgi,f6p}}$ | $v_{\mathrm{zwf}}$ | $K_{\mathrm{m,zwf,g6p}}$ | $v_{\mathrm{pfk}}$ | $K_{\mathrm{m,pfk,f6p}}$ | $n_{\mathrm{pfk}}$ |
|---|---|---|---|---|---|---|---|---|---|
| Value | 0.8751 | 0.3000 | 0.2800 | 0.1470 | 0.0853 | 0.1000 | 2.615 | 0.1600 | 3 |
| Ref. | est. using[54] | [84] | BRENDA (Gao2005) | BRENDA (Gao2005) | est. using[54] | BRENDA (Banerjee1972) | est. using[54] | BRENDA (Zheng1995) | [84] |
| Unit | mM/s | n.a. | mM | mM | mM/s | mM | mM/s | mM | n.a. |

| Param | $v_{\mathrm{ino1}}$ | $K_{\mathrm{m,ino1,g6p}}$ | $v_{\mathrm{t,mi}}$ | $K_{\mathrm{m,t,mi}}$ | $v_{\mathrm{miox}}$ | $K_{\mathrm{m,miox,mi}}$ | $a_{\mathrm{miox}}$ | $K_{\mathrm{a,miox,mi}}$ |
|---|---|---|---|---|---|---|---|---|
| Value | 0.2616 | 1.1800 | 0.0450 | 15 | 0.2201 | 24.7 | 5.4222 | 20 |
| Ref. | scaled val. from[53] | BRENDA (Donahue1981) | guess | guess | scaled val. from[83] | [86] | est. from data in[53] | est. from data in[53] |
| Unit | mM/s | mM | mM/s | mM | mM/s | mM | n.a. | mM |

The reaction rates and their assumed mechanisms are listed in Table S1. The value of each model parameter, reference of its source, and its units are shown in Table S2. Values are already scaled from those at the source so as to report them at the correct units of concentration (mM) and flux (mM/s). The kinetics of the MI-transport reaction $v_{\mathrm{t,m}}$ are unknown. We fixed values of its kinetic parameters $K_{\mathrm{m,t,mi}}$ and $v_{\mathrm{m,t,mi}}$ assuming competition for MI between $v_{\mathrm{t,m}}$ and $v_{\mathrm{Miox}}$ are of similar order, but where flux of transport is drastically smaller than consumption by MIOX. The $v_{\max}$ values of Ino1 and MIOX sourced from literature were converted from values $\widetilde{v}_{\mathrm{m,enz}}$ in units nmol/min/mg-Protein to values $v_{\mathrm{m,enz}}$ in units $\mathrm{mMs}^{-1}$ as follows:

$$v_{\mathrm{m,enz}} = \widetilde{v}_{\mathrm{m,enz}} \cdot \frac{\mathrm{TotCellProt}}{\mathrm{CellVol}} \cdot \frac{1}{60 \times 10^6}, \tag{S5}$$

where total protein content TotCellProt = $191 \times 10^{-12}$ mg-Protein/cell, in agreement with sources[87,88], and cell volume CellVol = $4.96 \times 10^{-16}$ L/cell[89]. The $v_{\max}$ values of the native enzymes were estimated using steady state data[54]. Given that reaction equations can be written in the form $f_{\text{enz}} = v_{\text{m,enz}} \cdot r_{\text{enz}}(m, p)$, their values were calculated as follows:

$$v_{\text{m,enz}} = \frac{f_{\text{ss,enz}}}{r_{\text{enz}}(m_{\text{ss}}, p)}, \tag{S6}$$

where $f_{\text{ss,enz}}$ is the steady state flux value of the respective reaction from fluxomics data and $m_{\text{ss}}$ is the corresponding steady state metabolite concentration from metabolomics data[54], shown in Table S3. These parameters are consistent with the growth rate and glucose uptake rate set in the model, as stated in Eq. (S1) and Eq. (S2).

TABLE S3. **Steady state multiomics data used to estimate $\mathbf{v}_{\max}$ values of native reactions.** All data are sourced from Keio multi-omics dataset[54], for continuous culture chemostat experiments with dilution rate 0.1h$^{-1}$. All fluxes are given as a percent of glucose uptake rate $v_{\text{PTS}}$ in Eq. (S2), and all concentrations are given in units of mM.

| Metabolites | | Fluxes | | |
|---|---|---|---|---|
| g6p | f6p | $f_{\text{Pgi}}$ | $f_{\text{Zwf}}$ | $f_{\text{PfkA}}$ |
| 0.281 | 0.0605 | $0.62 \cdot (v_{\text{PTS}} - \lambda \cdot g6p)$ | $0.38 \cdot (v_{\text{PTS}} - \lambda \cdot \text{g6p})$ | $f_{\text{Pgi}} + 0.5 \cdot f_{\text{Zwf}} - \lambda \cdot \text{f6p}$ |

### Dynamics of pathway enzymes

We assume that native enzymes are constitutively expressed, and so their concentrations remain constant in the timescale of the culture. Conversely, we model the dynamics of Ino1 and MIOX via Michael-Menten kinetics parameterised with $K_{\text{m}}$ and $v_{\text{max}}$ values from the literature. The maximum reaction rate of an enzyme $E$ is typically proportional to its concentration, so that $v_{\text{max}} = k_{\text{cat}} \times E$. But as we do not have $k_{\text{cat}}$ values for Ino1 and MIOX, we choose to model the maximum reaction rate as $v_{\text{max}} = \hat{v}_{\text{max}} \cdot E_{\text{rel}}$, Here, $\hat{v}_{\text{max}}$ is the value from literature and $E_{\text{rel}}$ is a non-dimensional variable representing the relative abundance of the enzyme. To this end, we model the dynamics of the relative abundance of Ino1 and MIOX with the ODEs:

$$\begin{aligned} \frac{\text{dIno1}_{\text{rel}}}{\text{d}t} &= u_{\text{ino1}}(\text{MI}) - \lambda \cdot \text{Ino1}_{\text{rel}}, \\ \frac{\text{dMIOX}_{\text{rel}}}{\text{d}t} &= u_{\text{miox}}(\text{MI}) - \lambda \cdot \text{MIOX}_{\text{rel}}. \end{aligned} \tag{S7}$$

The first term of each equation represents the expression rate and is modelled as either a constant (no control) or as a Hill function when expression is controlled by the biosensor IpsA through binding to *myo*-inositol (MI):

$$u_i(\text{MI}) = \begin{cases} a_i, & \text{no control}, \\ \dfrac{a_i \cdot \theta_i^{n_i}}{\theta_i^{n_i} + \text{MI}^{n_i}}, & \text{repression}, \\ \dfrac{a_i \cdot \text{MI}^{n_i}}{\theta_i^{n_i} + \text{MI}^{n_i}}, & \text{activation}, \end{cases} \quad i = \{\text{ino1, miox}\} \tag{S8}$$

where $\theta_i$ is the threshold concentration of MI to bring the expression rate halfway between its maximum and minimum and we fix the Hill coefficients to $n_{\text{ino1}} = n_{\text{miox}} = 1$. The $a_i$ parameters (in units of s$^{-1}$) are the maximal expression rate for the relative enzyme abundances.

### SUPPLEMENTARY FIGURES

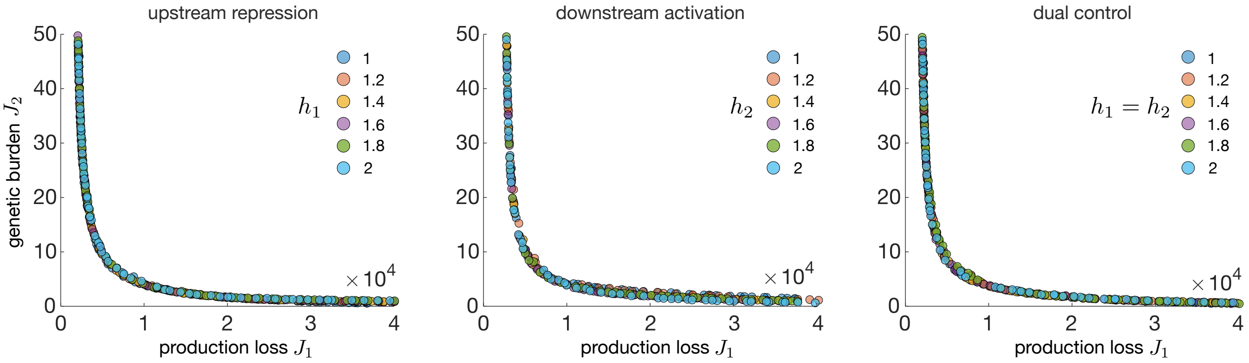


Supplementary Figure 1. **Impact of Hill coefficient on Pareto fronts.** Optimal solutions for the example pathway in Fig. 3A for various values of the Hill coefficient. The resulting Pareto fronts are largely insensitive to the Hill coefficient.

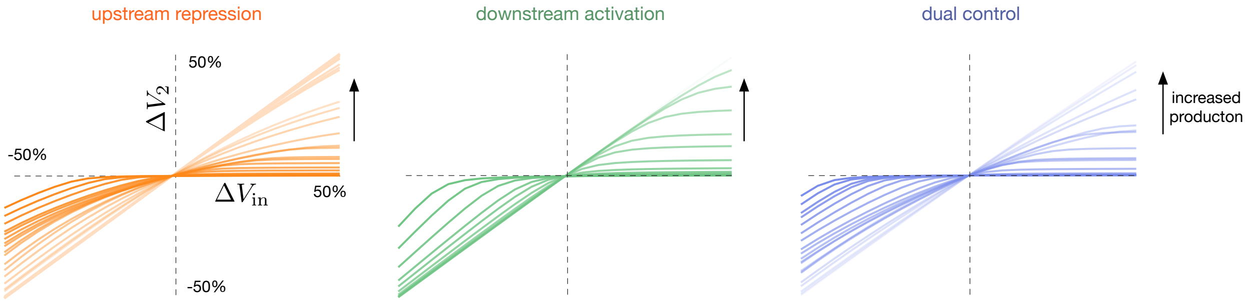


Supplementary Figure 2. **Response of optimal circuits to changes in growth conditions.** Response to perturbations in carbon influx for each optimized control circuit along the Pareto fronts in Fig. 3B. Each plot shows the relative change in steady state production flux $V_2$ for a given relative change in the carbon flux $V_{\text{in}}$; light (dark) coloured lines correspond to high (low) production regimes, i.e. the left (right) part of the Pareto front in Fig. 3B. The slopes of the curves represent the robustness of the production flux. The downstream activation architecture has a robust response for increases in carbon flux ($\Delta V_{\text{in}} > 0$), but not for reductions in carbon flux ($\Delta V_{\text{in}} < 0$), except at the low production regime. In contrast, the other two architectures display a robustness that is more evenly distributed across the Pareto front, as illustrated by Fig. 4B. In all cases, the steady state production flux $V_2$ was computed by first pre-simulating the models to a steady state with a constant $V_{\text{in}} = 1\mu\text{Ms}^{-1}$, and then introducing a step perturbation in $V_{\text{in}}$ until the dynamics reach a new steady state.

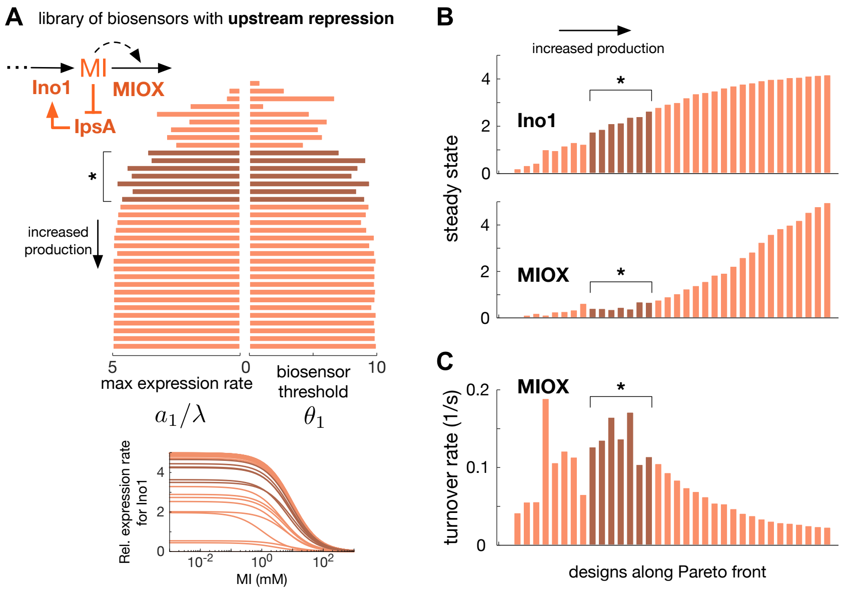


Supplementary Figure 3. **Optimal upstream repression circuits for production of glucaric acid.** **(A)** Optimal library of *myo*-inositol biosensors; bars are the maximal expression rates in dimensionless units ($a_1/\lambda$) and threshold ($\theta_1$) of the biosensor that controls Ino1 expression; the respective dose-response curves are shown in the inset below. **(B)** Steady state fold-change in Ino1 and MIOX enzyme abundance for each Pareto front solution in panel A. **(C)** Steady state turnover rate (flux per unit enzyme) of MIOX for each Pareto front solution. Dark shaded bars in all panels, and dose-response curves, represent the optimal solutions in the optimal trade-off region of the Pareto front, marked with an asterisk in Fig. 6A.

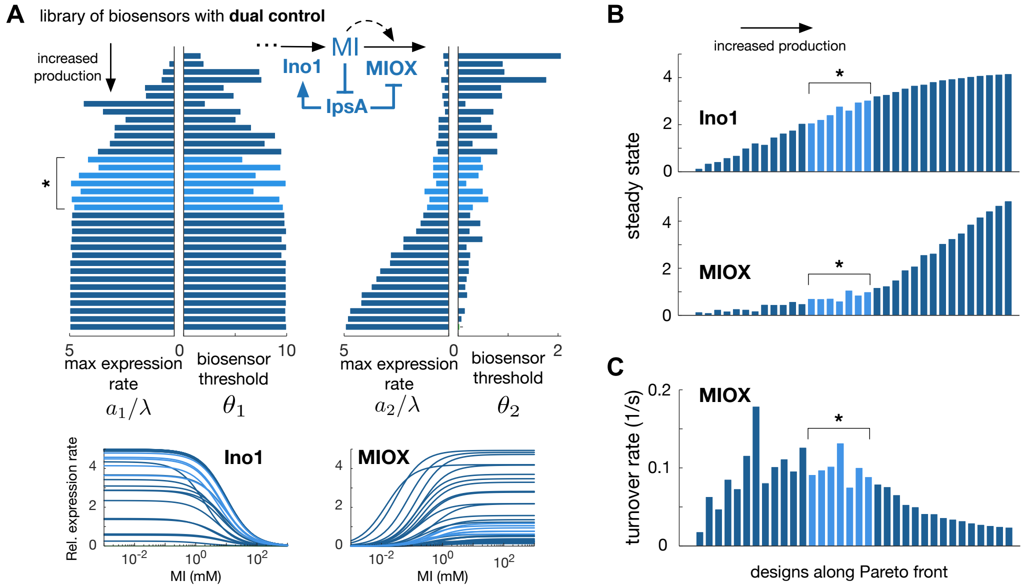


Supplementary Figure 4. **Optimal dual control circuits for production of glucaric acid.** **(A)** Optimal library of *myo*-inositol biosensors; bars are he maximal expression rates in dimensionless units ($a_i/\lambda$) and biosensor thresholds ($\theta_i$) that control Ino1 and MIOX; the respective dose-response curves are shown in the inset below. **(B)** Steady state fold-change in Ino1 and MIOX enzyme abundance for each Pareto front solution in panel A. **(C)** MIOX steady state turnover rate (flux per unit enzyme) for each Pareto front solution. Light shaded bars in all panels and dose-response curves represent the optimal solutions in the optimal trade-off region of the Pareto front, marked with an asterisk in Fig. 6A.
